# CMAC: A deep learning framework for absolute cardiomyocyte transcriptional age estimation

**DOI:** 10.64898/2026.09.19.752897

**Authors:** Kwaku K. Quansah, Nadine Zureick, Elaine Chen, Sean Murphy, Emma Anderson, Esther Kwon, David Suh, Genevieve Stein-O’Brien, Patrick Cahan, Chulan Kwon

## Abstract

Cardiomyocyte maturation spans embryogenesis through postnatal life and involves coordinated transcriptional transitions that are disrupted in disease and incompletely recapitulated by stem cell-derived cardiomyocytes. However, no unified framework exists for assigning absolute transcriptional maturation age across datasets, platforms, and experimental conditions. Here, we developed CMAC (Cardiomyocyte Maturation Age Clock), integrating 18 murine single-cell RNA-sequencing datasets spanning embryonic and postnatal timepoints, into a batch-invariant probabilistic atlas using single cell variational inference (scVI) with a 100-dimensional latent representation. Analysis of the developmental trajectory revealed highly non-uniform transcriptional change, with 63.9% of the total transcriptional journey completed by birth and the neonatal transition representing the largest single transcriptional step. An XGBoost regressor trained on the CMAC latent space predicted absolute chronological age with a leave-one-dataset-out mean absolute error (MAE) of 4.09 days (median 3.27 days), Pearson *r* = 0.903, and Spearman ρ = 0.908 across 18 datasets and six sequencing platforms, outperforming linear regression (MAE = 6.18 days) and PCA-based representations (MAE = 6.35 days). In an independent P14 cohort comprising 98,163 cardiomyocytes from six biological replicates, CMAC achieved a mouse-level MAE of 0.33 days, with all six mice predicted within 1.1 days of chronological age. Application of CMAC to a previously published PGC1α/β double-knockout model with established delayed cardiomyocyte maturation independently recapitulated this phenotype, quantifying maturation deficits of 4.77 to 8.60 days across P7 to P28, with the largest deficit at P14. Together, these results establish CMAC as a quantitative, deep learning framework for assigning absolute transcriptional maturation age and quantifying deviations from normal cardiomyocyte development.

## Introduction

Cardiomyocyte maturation is the last phase of heart development, in which fetal cardiomyocytes progressively acquire the structural, metabolic, electrophysiological, and contractile properties required for post-birth cardiac function^1^. Underlying this process is a complex, non-linear transcriptional program that is frequently disrupted in cardiomyopathies.^2–4^ Efforts to generate pluripotent stem cell (PSC)-derived cardiomyocytes for biomedical applications are hindered by the persistent transcriptional immaturity of these cells relative to their in vivo counterparts.^5,6^ Accurately quantifying cardiomyocyte maturation is therefore essential for studying cardiac development, interpreting disease phenotypes, and evaluating PSC-derived cardiomyocyte models.

Cardiomyocyte maturity has traditionally been evaluated using combinations of morphological, electrophysiological, metabolic, contractile, and molecular features.^7^ Contemporary transcriptome-based approaches have provided more scalable quantitative measures.^8,9^ Bulk and microarray-based methods such as MatStat compare sample-level gene expression to developmental gene regulatory networks but cannot resolve single-cell heterogeneity.^10^ Relative-expression-ordering approaches have scored PSC-derived cardiomyocytes according to the fraction of gene-pair relationships resembling adult myocardium, while other methods derive maturation scores from principal components of developmental cardiac transcriptomes.^11^ Pseudotime inference methods including Monocle have been applied to order cardiomyocytes along inferred developmental trajectories and detect maturation deficits in genetic perturbation models.^12^ More recently, Kannan et al. developed a single-cell transcriptomic entropy score that uses gene distribution to estimate maturation state at the single-cell level, subsequently formalized as a cross-study workflow for quantifying PSC-derived cardiomyocyte maturation from scRNA-seq data.^13^ Despite these advances, existing maturation metrics share two fundamental limitations. First, they provide relative measures of maturity rather than assigning cardiomyocytes an absolute developmental age in interpretable units, making direct cross-study comparison difficult. Second, integration of developmental data across studies remains challenging because datasets differ in cell isolation protocol, library preparation chemistry, sequencing platform, developmental timepoint sampling, and sequencing depth, introducing technical variation that confounds biological comparisons.^14,15^ A framework that learns a shared developmental representation across heterogeneous datasets and translates that representation directly into chronological age would provide an intuitive common scale: for example, determining whether a cardiomyocyte from a disease model or differentiation protocol transcriptionally resembles a typical P7, P14, or P21 wild-type cardiomyocyte.

Here, we developed the Cardiomyocyte Maturation Age Clock (CMAC) by integrating 18 murine single-cell RNA-sequencing datasets spanning embryonic day 7.75 (E7.75) to postnatal day 28 (P28) using a deep generative model with per-dataset batch correction. Single cell variational inference (scVI) was selected for its ability to model raw count data using a negative binomial likelihood and to explicitly separate technical variation attributable to sequencing platform and laboratory of origin from biological developmental signal through per-dataset batch embeddings.^16^ An XGBoost regressor was then trained to predict chronological developmental age from the resulting latent representation.^17^ We show that transcriptional maturation proceeds non-uniformly across development, with 63.9% of the total transcriptional journey complete at birth and the neonatal transition representing the largest single transcriptional step. We evaluate CMAC using leave-one-dataset-out (LOGO) cross-validation across 18 datasets and six sequencing platforms, and validate generalization to a truly held-out cohort of 98,163 P14 cardiomyocytes from six biological replicates. Finally, we apply CMAC to cardiomyocytes from PGC1α/β double-knockout mice, in which impaired postnatal maturation has been previously characterized, demonstrating that CMAC detects and quantifies this known phenotype as a deficit in predicted developmental age. CMAC thus provides a transferable framework for placing cardiomyocytes from diverse experimental contexts onto a common, quantitatively interpretable developmental timescale.

## Results

### CMAC learns a compressed representation of cardiomyocyte maturation

Building a unified reference for cardiomyocyte maturation requires integrating transcriptomic data collected across different laboratories, sequencing platforms, and experimental designs without allowing technical variation to obscure genuine developmental biology. We assembled 18 murine scRNA-seq datasets spanning embryonic day 7.75 (E7.75) through postnatal day 28 (P28), encompassing 8,294 cardiomyocytes across six sequencing platforms, **(Supplemental Table 1)**.^18–27^ To ensure that selected genes reflect genuine developmental variation rather than dataset-specific technical effects, highly variable genes were identified by computing between-stage variance on a balanced pseudobulk representation that gives each dataset equal weight at each developmental stage, yielding 2,991 maturation-variable genes after exclusion of mitochondrial, ribosomal, and pseudogene features.

We selected scVI, a variational autoencoder, as the integration framework for three reasons. First, scVI models raw count data using a negative binomial likelihood, which explicitly accounts for the overdispersion and zero-inflation characteristic of scRNA-seq count matrices, providing a more principled generative model than methods that operate on normalized or log-transformed expression. Second, scVI’s per-dataset batch embedding separates technical variation from biological variation in the latent space, enabling cells from different experimental systems to be compared on a common developmental coordinate. Third, the learned latent representation supports reference mapping via scVI’s load_query_data functionality, allowing new datasets to be projected into the trained atlas without retraining, which is essential for CMAC’s application as a transferable scoring framework.^28^ Together these properties make scVI well-suited for constructing a developmental atlas that must generalize across the diverse technical backgrounds of 18 independently generated datasets.^16^

Thus, we applied scVI to learn a 100-dimensional batch-corrected latent representation, assigning each dataset a unique batch label to account for platform and laboratory effects. Three datasets were excluded following preliminary leave-one-dataset-out evaluation due to catastrophic misprediction attributable to fixation protocol artifacts or chamber-specific transcriptional identity. The learned latent space organized cells continuously by developmental stage, with early embryonic progenitors and late postnatal cardiomyocytes occupying distinct but continuously connected regions of the embedding **(Figure 1A)**. Dataset labels were uniformly distributed throughout the embedding with no single dataset dominating any region, confirming successful batch correction across six sequencing platforms **(Figure 1B)**. Developmental timepoints were similarly well-distributed, with smooth transitions between adjacent stages visible in the categorical timepoint embedding **(Figure 1C)**. Integration quality was confirmed quantitatively by a dataset silhouette score of −0.076, cross-dataset neighbor fraction of 55.4%, iLISI median of 2.527, age-neighborhood Spearman correlation of 0.893, and median neighbor age difference of 2.77 days, together demonstrating that scVI successfully isolated biological developmental variation from technical batch effects **(Figure 1D)**.

**Figure 1.**
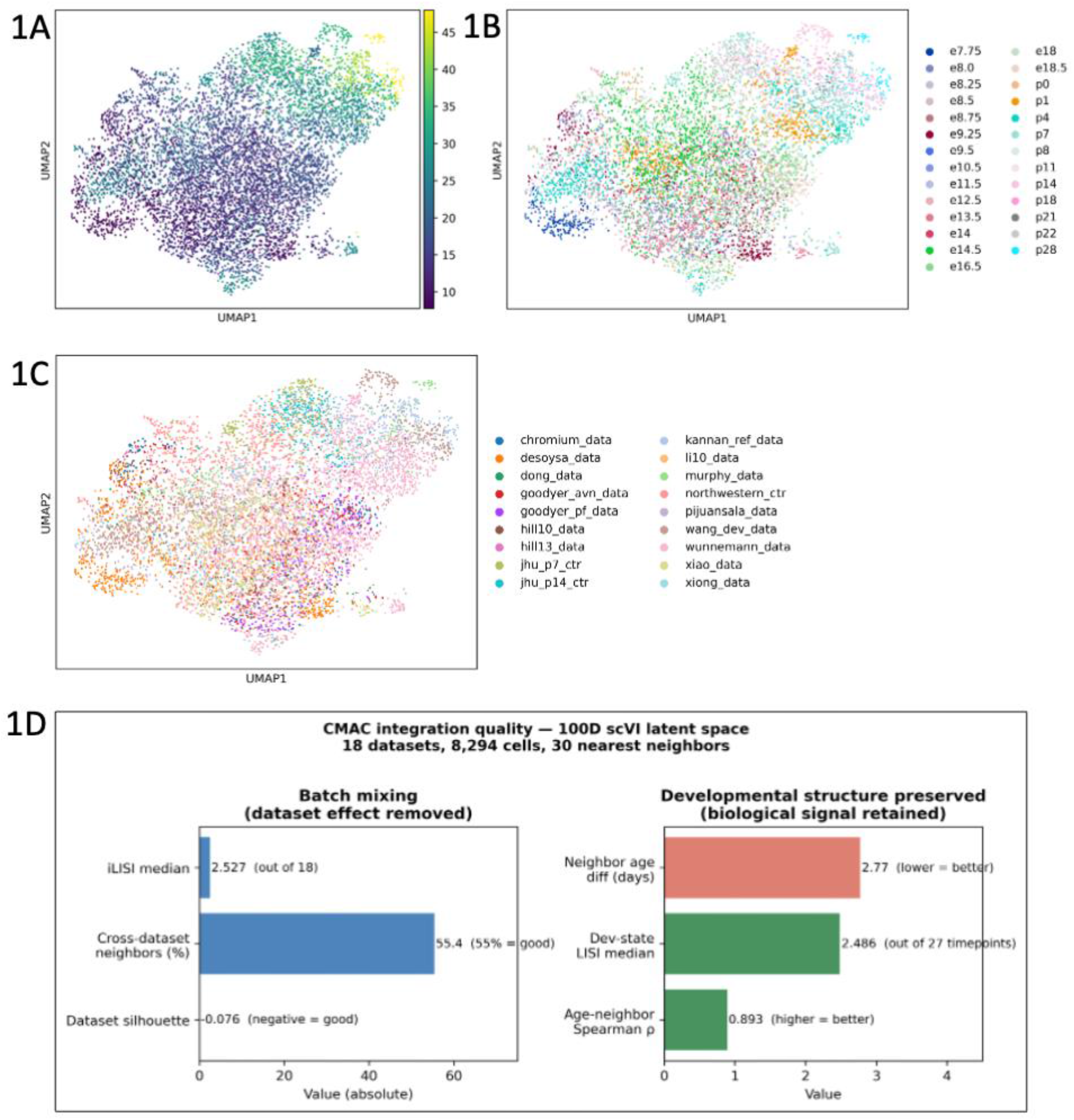
CMAC learns a compressed representation of cardiomyocyte maturation. **1A** UMAP visualization of the 100-dimensional scVI latent space colored by biological age (days post conception), revealing continuous developmental organization from early embryonic progenitors (dark) through late postnatal cardiomyocytes (light). **1B** The same UMAP colored by dataset of origin, demonstrating successful batch correction across 18 independent datasets and six sequencing platforms with no single dataset dominating any region of the embedding. **1C** The same UMAP colored by developmental timepoint. **1D** Integration quality metrics computed from the scVI latent space (30 nearest neighbors): dataset silhouette score (−0.076, negative values indicate cross-dataset mixing), cross-dataset neighbor fraction (55.4%), iLISI median (2.527), age-neighborhood Spearman correlation (0.893), and median neighbor age difference (2.77 days), confirming successful removal of technical batch effects while preserving developmental age as the primary organizer of the latent space.

### CMAC reconstructs a continuous cardiomyocyte maturation manifold

To demonstrate that the CMAC latent space captures genuine biological structure rather than technical variation, we examined the spatial organization of known maturation markers within the embedding.^29^ The fetal contractile isoforms *Myh6* (β-myosin heavy chain) *Tnni1* (slow skeletal troponin I) were highly expressed throughout the embryonic body of the UMAP and decreased progressively in the postnatal arm, while the adult isoforms Myh7 and Tnni3 showed the reciprocal pattern, emerging in the postnatal arm consistent with the known isoform switching program of cardiomyocyte maturation **(Figure 2A)**.

**Figure 2.**
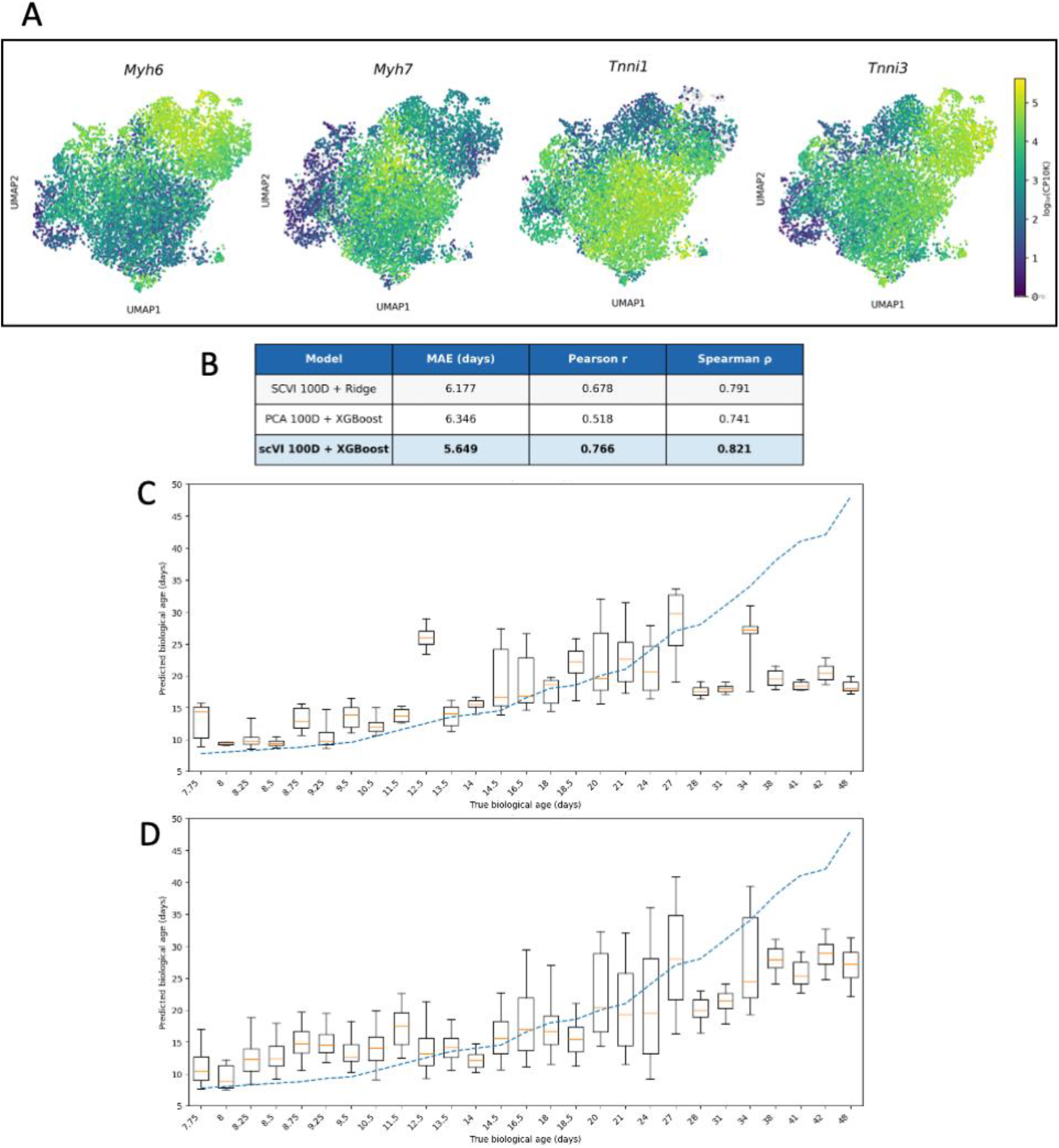
CMAC reconstructs a continuous cardiomyocyte maturation manifold. **2A**. Expression of canonical cardiomyocyte maturation markers overlaid on the CMAC UMAP. *Myh6* (fetal myosin heavy chain) and *Tnni1* (fetal troponin I) are highly expressed in the embryonic body and decrease postnatally, while *Myh7* (adult myosin heavy chain) and *Tnni3* (adult troponin I) emerge in the postnatal arm, confirming that the latent space captures the known isoform switching program of cardiomyocyte maturation. Expression shown as log_10_(CP10K). **2B** Comparison of prediction distributions for scVI 100D + XGBoost versus PCA 100D + XGBoost across all developmental timepoints. Each box represents the distribution of predicted ages for cells at a given true age. scVI integration substantially outperforms PCA across all metrics (scVI: MAE = 5.65 days, r = 0.77, ρ = 0.82; PCA: MAE = 6.35 days, r = 0.52, ρ = 0.74), demonstrating that explicit batch correction and nonlinear dimensionality reduction are essential for recovering developmental age signal across 18 independent datasets.

To demonstrate that the non-linear generative model captures developmental structure beyond what simpler methods achieve, we compared CMAC’s scVI latent space to a 100-dimensional PCA decomposition of the same log-normalized count matrix paired with the same XGBoost regressor.^16,17^ The prediction distributions per timepoint were substantially tighter and better aligned with the identity line for scVI than for PCA, particularly in the late postnatal window where PCA predictions collapsed toward a narrow range regardless of true age **(Figure 2B)**. Across all quantitative metrics scVI outperformed PCA (MAE: 5.65 vs 6.35 days; Pearson r: 0.77 vs 0.52; Spearman ρ: 0.82 vs 0.74), demonstrating that explicit batch correction and non-linear dimensionality reduction are essential for recovering developmental age signal across 18 independent datasets. scVI similarly outperformed a Ridge regression baseline trained on the same latent space (MAE: 5.65 vs 6.18 days; Pearson r: 0.77 vs 0.68), confirming that non-linear XGBoost regression captures developmental structure beyond what a linear model can extract from the latent representation **(Figure 2B)**.

### Transcriptional change is non-uniform across cardiomyocyte development

To characterize the structure of the developmental trajectory, we computed the cosine distance between adjacent timepoint mean latent vectors, measuring the directional change in transcriptional program between each consecutive developmental stage. Cumulative summation of these distances from E7.75 to P28, normalized to a 0 to 1 scale, revealed that transcriptional change is highly non-uniform across cardiomyocyte development. Plotting cumulative transcriptional distance against chronological age produced a pronounced S-shaped curve, deviating substantially from the linear expectation that equal chronological time would correspond to equal transcriptional change **(Figure 3A)**. The largest single transition occurred at birth (P0, cosine distance = 1.614), consistent with the dramatic metabolic and structural reorganization of cardiomyocytes at the neonatal transition, and 63.9% of the cumulative transcriptional journey was complete at birth despite the entire postnatal window spanning more absolute days than embryonic development^30,31^ **(Figure 3A)**. The late postnatal window from P14 to P28 contributed only 8.6% of total transcriptional change despite spanning 14 chronological days **(Figure 3A)**.

**Figure 3.**
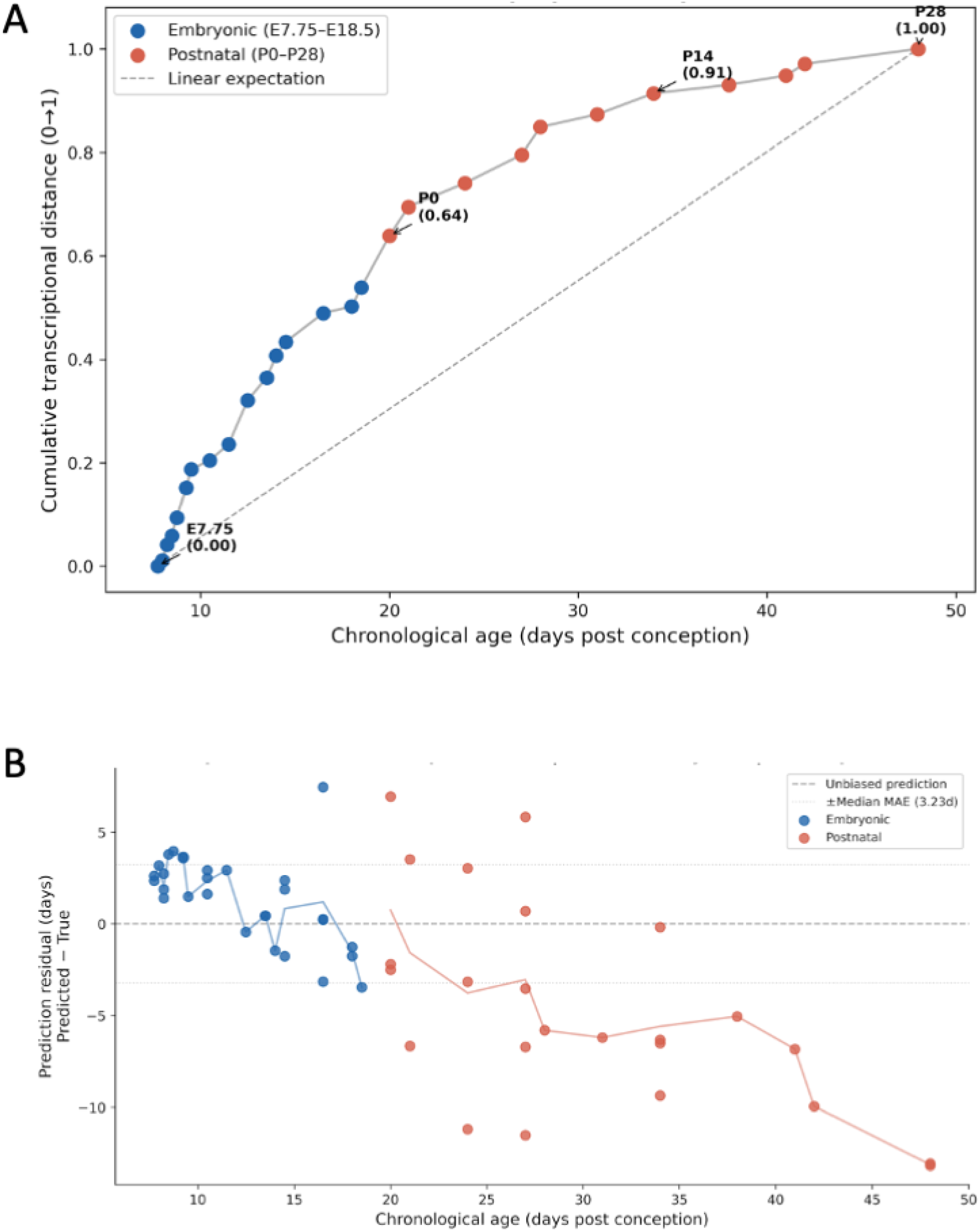
Transcriptional change is non-uniform across cardiomyocyte development. **3A** Cumulative transcriptional distance (normalized cosine distance along the developmental trajectory, 0-1) plotted against chronological age (days post conception) for each developmental timepoint. The S-shaped curve reveals that transcriptional change is concentrated in the embryonic and early neonatal period. 63.9% of the total transcriptional journey is complete at birth (P0), while late postnatal stages (P14–P28) contribute relatively little additional transcriptional change. The dashed line shows the linear expectation if chronological age and transcriptional change were proportional. Key developmental milestones are annotated. **3B** LOGO cross-validation residuals (predicted minus true age) per dataset × timepoint group, colored by developmental phase (blue = embryonic, red = postnatal). Each point represents the mean residual for one dataset × timepoint combination. Embryonic timepoints show slight positive bias (+2 to +4 days) while postnatal timepoints show increasing negative bias, consistent with the slower rate of transcriptional change in the postnatal window. The dotted lines indicate the CMAC median prediction error (±3.27 days).

This non-uniformity has direct implications for interpreting maturation deficits: equal chronological time does not represent equal transcriptional progression, and a given prediction error carries different biological significance depending on developmental stage. Residual analysis of LOGO cross-validation predictions confirmed this pattern. Embryonic timepoints showed slight positive bias (+2 to +4 days) while postnatal timepoints showed increasing negative bias consistent with slower transcriptional change per unit time, with the dotted reference lines **(Figure 3B)** indicating that most embryonic timepoints fall within the CMAC median prediction error of ±3.27 days while several postnatal datasets exceed this threshold **(Figure 3B)**.

### CMAC predicts absolute cardiomyocyte maturation age across platforms and datasets

We trained an XGBoost regressor on the CMAC latent space to predict chronological age in days post-conception, using log-transformed age as the target and combined dataset × stage inverse-frequency weighting to address the severe imbalance in cell counts across developmental stages. Hyperparameters were selected via nested cross-validation using 53 dataset × developmental stage groups, yielding consensus parameters that emphasized shallow trees (max_depth=3) with moderate regularization, consistent with the relatively small training set size relative to the 100-dimensional feature space.

The cumulative transcriptional distance trajectory confirmed the biological calibration of the latent space, with birth representing the single largest transcriptional step (cosine distance = 1.614) and 63.9% of the total transcriptional journey complete at birth, while the late postnatal window from P14 to P28 contributed only 8.6% of total change **(Figure 4A-B)**. Leave-one-dataset-out cross-validation across all 18 datasets yielded a mean absolute error of 4.09 days (median 3.27 days), Pearson r = 0.903, and Spearman ρ = 0.908 at the dataset × developmental stage level, with predicted ages tracking true ages closely across the full E7.75 to P28 developmental window **(Figure 4C)**. Performance was strongest for multi-timepoint datasets where both MAE and rank correlation could be assessed, while single-timepoint datasets showed a range of MAEs reflecting platform-specific batch effects of varying severity **(Figure 4D)**.

**Figure 4.**
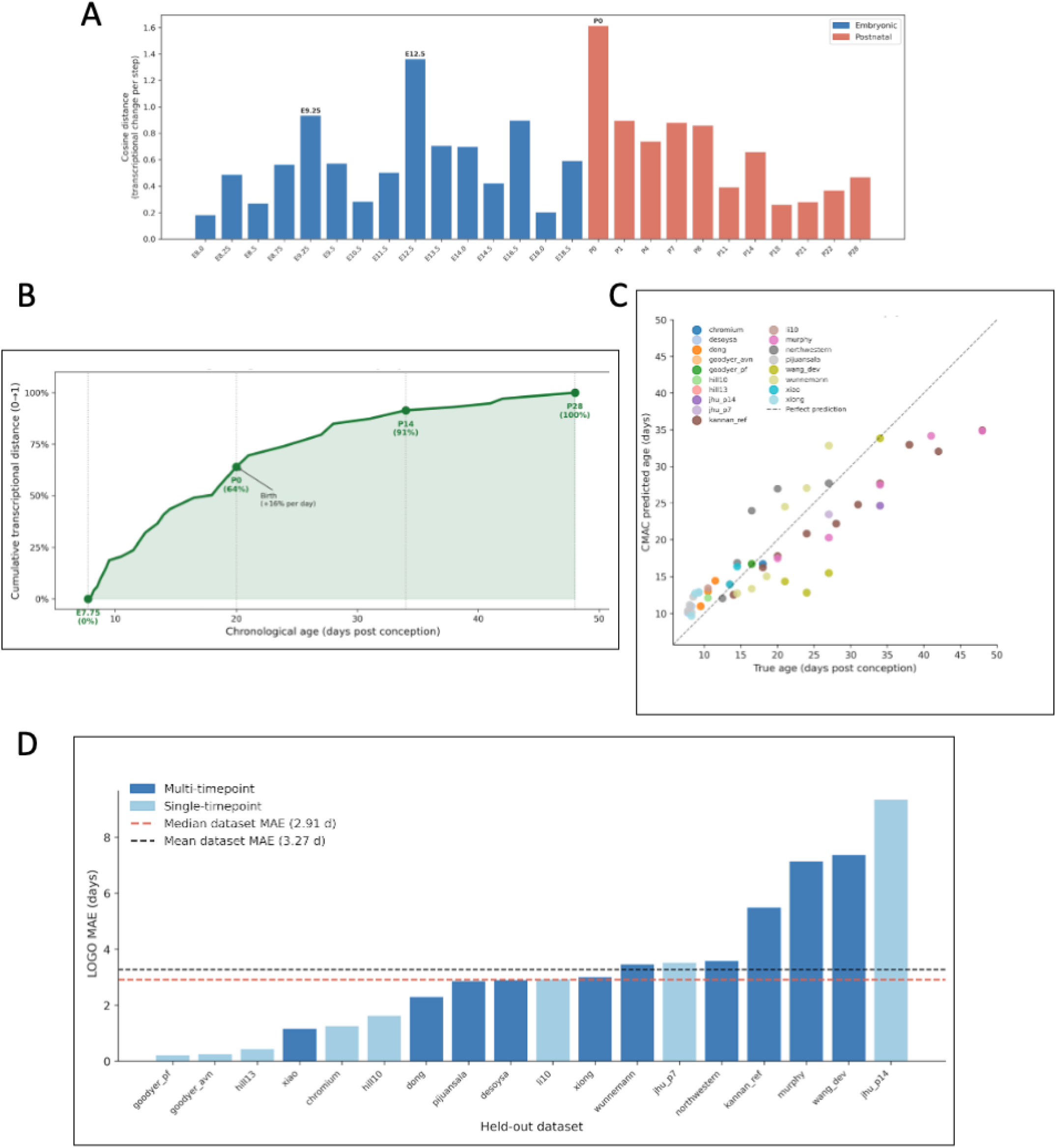
CMAC predicts absolute cardiomyocyte maturation age across platforms and datasets. **4A** Cosine distance between adjacent developmental timepoint centroids in the CMAC latent space, quantifying the transcriptional change contributed by each developmental transition from E7.75 to P28. The largest single transition occurs at birth (P0, cosine distance = 1.614), consistent with the dramatic metabolic and structural reorganization of cardiomyocytes at the neonatal transition. Blue bars indicate embryonic transitions; red bars indicate postnatal transitions. **4B** Cumulative transcriptional distance along the developmental trajectory normalized to a 0 to 1 biological age scale, where 0 represents the earliest cardiac progenitor state (E7.75) and 1.0 represents completed postnatal maturation (P28). Equal increments on this scale correspond to equal amounts of transcriptional change. Key developmental milestones are annotated with their cumulative score and chronological age, highlighting that 63.9% of the total transcriptional journey is complete at birth and 91.4% by P14. **4C** Leave-one-dataset-out cross-validation scatter plot showing mean predicted age versus true age for each of 53 (dataset × developmental stage) groups across all 18 held-out datasets. Each color represents one held-out dataset. The dashed line indicates perfect prediction. Overall performance: MAE = 4.09 days, Pearson r = 0.903, Spearman ρ = 0.908. **4D** Per-dataset LOGO mean absolute error sorted from lowest to highest. Dark blue bars indicate multi-timepoint datasets for which both MAE and rank correlation are computable; light blue bars indicate single-timepoint datasets for which only MAE is available. The red dashed line indicates the median MAE (3.27 days) and the black dashed line indicates the mean MAE (4.09 days) across all 18 datasets.

### CMAC generalizes to held-out data and detects disease-relevant maturation deficits

To first establish that CMAC generalizes accurately to held-out biological replicates at the most clinically relevant maturation timepoint, we applied the framework to an independent cohort of 98,163 P14 wild-type cardiomyocytes from 6 mice not used in any stage of model development. CMAC predicted a mean age of 33.89 ± 0.54 days across animals (true age 34.0 days), with a mouse-level MAE of 0.33 days and cell-level MAE of 3.16 days **(Figure 5A)**. All 6 mice were predicted within ±1.1 days of true age, with prediction errors ranging from −1.09 to +0.55 days across animals **(Figure 5B)**, demonstrating precise generalization to biological replicates from a truly held-out cohort.

**Figure 5.**
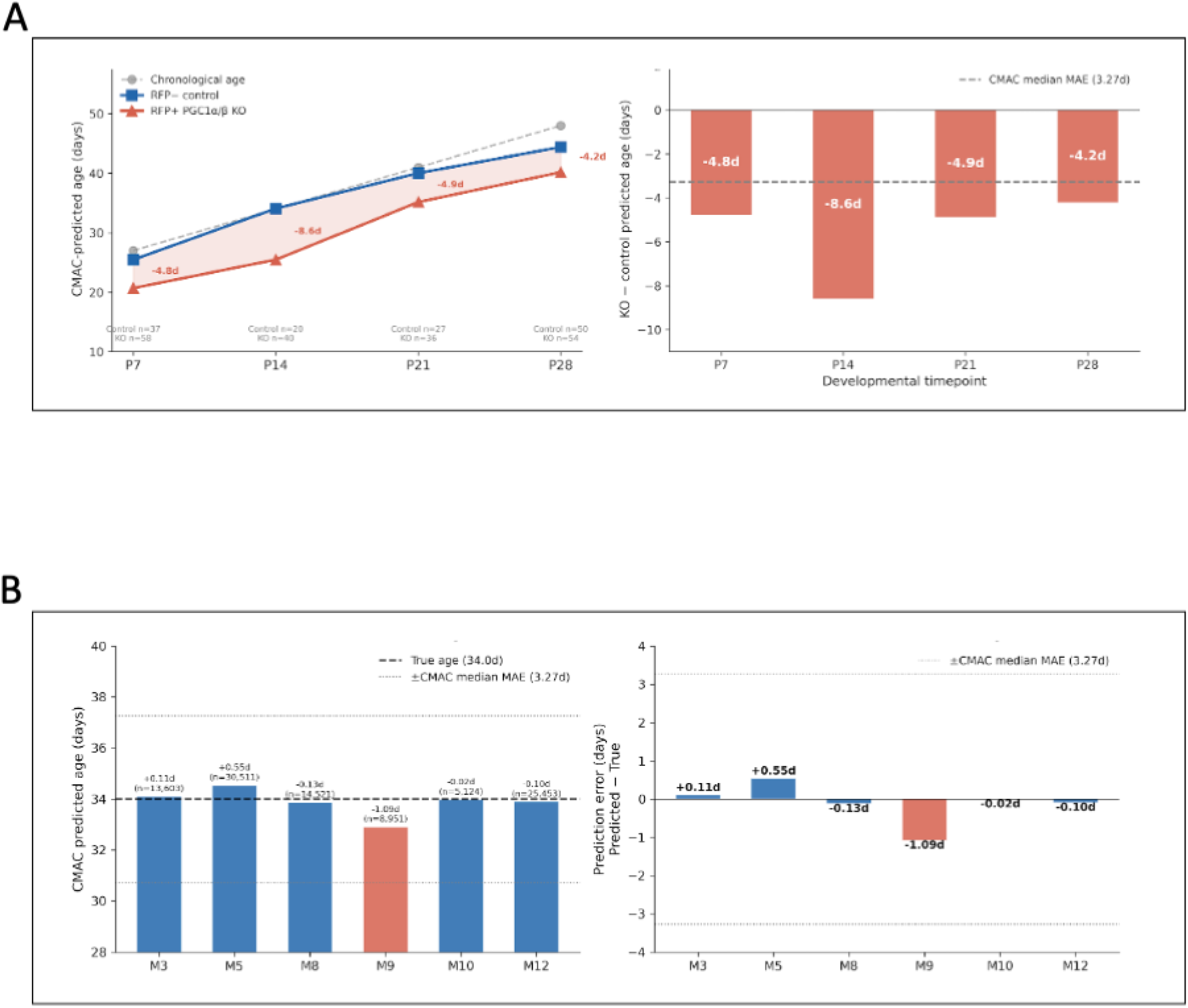
CMAC generalizes to held-out data and detects disease-relevant maturation deficits. **5A** External validation on an independent held-out P14 cohort of 98,163 cardiomyocytes from 6 biological replicates not used in any stage of model development. Left: CMAC-predicted maturation age per mouse. Bars are colored blue if the per-mouse error is within ±1.0 day of true age (34.0 days, black dashed line); dotted lines indicate the CMAC median prediction error (±3.27 days). Right: per-mouse prediction error (predicted minus true age). All 6 mice were predicted within ±1.1 days of true age (mouse-level MAE = 0.33 days). **5B** Application of CMAC to PGC1α/β double-knockout cardiomyocytes. Left: CMAC-predicted maturation age for RFP^+^ PGC1α/β knockout and RFP^−^ wild-type control cardiomyocytes at postnatal days 7, 14, 21, and 28, alongside true chronological age (gray dashed line). KO cardiomyocytes consistently show lower predicted maturation age than wild-type controls, with the deficit peaking at P14 (−8.60 days). Cell counts per group are indicated below the x-axis. Right: maturation deficit (KO median minus control median predicted age) at each timepoint. The gray dashed line indicates the CMAC median cross-validation error (3.27 days); all deficits exceed this threshold. Effect sizes are reported without significance testing as individual cell-level replication structure could not be confirmed for this dataset.

Having established CMAC’s accuracy on held-out wild-type data, we next applied the framework to cardiomyocytes from PGC1α/β double-knockout mice, in which loss of the master regulators of mitochondrial biogenesis is expected to impair postnatal metabolic maturation.^32^ Knockout cells were identified by RFP expression (RFP^+^) and projected into the CMAC latent space via reference atlas mapping, with RFP^−^ cells serving as wild-type controls. Chronological age was predicted using the trained XGBoost model without any parameter adjustment or retraining.

CMAC predicted a consistent maturation deficit in PGC1α/β knockout cardiomyocytes relative to wild-type controls across all four examined timepoints, with the deficit trajectory clearly separated from both wild-type predictions and true chronological age at every stage **(Figure 5C)**. The predicted deficits were −4.77 days at P7, −8.60 days at P14, −4.87 days at P21, and −4.22 days at P28, peaking at P14 consistent with the established role of PGC1α/β coactivators in driving the glycolytic-to-oxidative phosphorylation metabolic switch that defines the second postnatal week of cardiomyocyte maturation^32,33^ **(Figure 5D)**. Wild-type control cells were predicted at ages closely matching their true chronological ages (mean offset ≤1.2 days at P7, P14, and P21), confirming that the observed deficits reflect genuine transcriptional immaturity in knockout cells rather than systematic model bias **(Figure 5C)**. All four deficits exceeded the CMAC median cross-validation error of 3.27 days shown as a reference line in Figure 5D, providing a model-calibrated threshold above which predictions reflect biological signal rather than prediction uncertainty. As individual cell-level replication structure could not be confirmed for this dataset, effect sizes are reported without significance testing.

## Discussion

Quantifying cardiomyocyte maturation state has remained a fundamental challenge in cardiac biology. Existing approaches including pseudotime inference, relative maturity scores, and entropy-based metrics are dataset-specific, expressed in arbitrary units, and platform-dependent, making cross-experiment comparison impossible without reanalysis. Here we present CMAC, a deep learning framework that addresses these limitations by integrating 18 independent murine scRNA-seq datasets into a unified latent space and training a supervised regressor to assign absolute chronological age in days post-conception to individual cardiomyocytes. CMAC achieves leave-one-dataset-out cross-validation MAE of 4.09 days and Pearson r = 0.903 across six sequencing platforms, and generalizes with mouse-level MAE of 0.33 days to a truly held-out cohort of 98,163 P14 cardiomyocytes from six biological replicates. Unlike entropy-based approaches that provide a relative measure of maturation state independent of developmental stage, CMAC assigns absolute chronological age in interpretable units directly comparable across studies, providing an intuitive common scale for placing cardiomyocytes from diverse experimental contexts onto a shared developmental timeline.

A key finding emerging from the CMAC trajectory analysis is that transcriptional change is profoundly non-uniform across cardiomyocyte development. The birth transition contributes the single largest step in transcriptional distance (cosine distance = 1.614), consistent with the dramatic metabolic and structural reorganization that accompanies the neonatal cardiopulmonary transition.^34^ By postnatal day 14, 91.4% of the cumulative transcriptional journey from cardiac progenitor to mature cardiomyocyte is complete, while the subsequent two weeks to P28 contribute only 8.6% of total transcriptional change. This non-uniformity has direct practical implications for interpreting maturation deficits: a deficit of a given number of chronological days carries fundamentally different biological significance depending on the developmental stage at which it occurs. This also explains why relative maturity scores and pseudotime metrics can misrepresent maturation deficits. For example, a deficit of equal pseudotime magnitude at P14 versus P21 represents a fundamentally different fraction of the remaining maturation program, a distinction that only an absolute, trajectory-aware metric can capture. The choice of P28 as the maturation endpoint reflects evidence that the major structural, metabolic, and electrophysiological transitions of cardiomyocyte maturation are largely complete by the fourth postnatal week.^19^

Application of CMAC to PGC1α/β double-knockout cardiomyocytes demonstrates its utility for detecting maturation deficits in disease-relevant genetic perturbation models. The predicted deficit peaked at P14 (−8.60 days), consistent with the established role of PGC1α/β coactivators in driving the glycolytic-to-oxidative phosphorylation metabolic switch that defines the second postnatal week of cardiomyocyte maturation.^35^ The deficit exceeded the CMAC median cross-validation error of 3.27 days at all four timepoints examined, providing a model-calibrated threshold for distinguishing biological signal from prediction uncertainty. Wild-type control cells were predicted at ages closely matching their true chronological ages, confirming that the observed deficits reflect genuine transcriptional immaturity in knockout cells rather than systematic model bias. We note that individual cell-level replication structure could not be confirmed for this dataset and effect sizes are therefore reported without significance testing; future applications of CMAC to datasets with confirmed biological replicates will enable formal statistical inference at the animal level.

Several limitations of the current implementation warrant discussion. First, the iLISI median of 2.527 out of a maximum of 18 indicates moderate rather than complete dataset mixing in the latent space. This reflects a structural constraint of the atlas rather than a failure of batch correction: many developmental timepoints are covered by only one or two datasets, making cross-dataset mixing at those timepoints structurally impossible regardless of the integration method. Among timepoints covered by two or more datasets, integration quality was substantially higher, and the age-neighborhood Spearman correlation of 0.893 confirms that the latent space is organized primarily by developmental age rather than dataset of origin. Nonetheless, additional datasets at underrepresented timepoints, where current cell counts fall below 75, would improve both integration quality and prediction accuracy. Planned Parse SPLiT-seq data at P7, P14, P21, and P28 from sex-balanced C57BL/6 cohorts will reinforce coverage at these timepoints and will be incorporated in a revised version of the atlas.

Second, one dataset (jhu_p14_ctr) showed systematic underprediction with a mean residual of −9.36 days, the largest of any dataset in the atlas. The contrasting performance of the independent Zureick_P14 held-out cohort (mouse-level MAE = 0.33 days), generated using a similar in-house protocol, suggests that this underprediction reflects batch-specific rather than protocol-level effects. Identifying and correcting for the source of this batch effect, which may relate to differences in library preparation run, sequencing depth, or reagent lot, remains an open question. When a small reference set of cells with known maturation state is available, a simple dataset-level calibration correction could substantially reduce this offset, analogous to calibration approaches used in epigenetic clock applications.

Third, CMAC is trained exclusively on murine scRNA-seq data spanning E7.75 to P28, and direct application to human cardiomyocytes or hPSC-derived cardiomyocytes requires cross-species adaptation that has not yet been validated. Extension of CMAC to human developmental atlases is a natural next step that would enable maturation benchmarking of hPSC-CM differentiation protocols, a pressing need given the known transcriptional immaturity of stem cell-derived cardiomyocytes relative to adult tissue.^5,36^ Fourth, the current atlas is restricted to scRNA-seq data; integration of snRNA-seq datasets, which are increasingly common for postnatal and adult cardiomyocytes due to their large cell size, will require additional modality-aware batch correction strategies not implemented in the current framework.

CMAC establishes a foundation for a broader computational framework for cardiomyocyte biology across the lifespan. The CMAC framework, integrating diverse public datasets into a reference atlas, learning a batch-invariant latent representation, and enabling transferable age prediction via reference mapping, provides a template for constructing developmental clocks in other cell types and organisms where maturation state is biologically and clinically relevant.

## Methods

### Dataset assembly

#### Publicly available datasets

15 publicly available and 3 in-house murine single-cell RNA-sequencing datasets were assembled to construct the Cardiomyocyte Maturation Age Clock (CMAC) **(Supplemental Table 1)**. Datasets were selected to provide broad coverage of cardiomyocyte development, spanning early embryonic stages through postnatal maturation and incorporating multiple sequencing platforms and experimental studies. Only *in vivo* murine cardiac datasets with UMI-based gene-expression measurements and identifiable developmental timepoints were included.

#### In-house dataset generation

Three datasets included in the CMAC atlas (jhu_p7_ctr, jhu_p14_ctr, northwestern_ctr) and one external validation dataset (Zureick_P14) were generated in-house and have not been previously published. All animal procedures were approved by the Johns Hopkins University Institutional Animal Care and Use Committee and performed in accordance with the NIH Guide for the Care and Use of Laboratory Animals.

#### Cardiomyocyte isolation

Cardiomyocytes were isolated using the Langendorff-free method of Ackers-Johnson et al. with minor modifications^37^. Mice were briefly anesthetized with isoflurane until loss of pedal reflex and euthanized by cervical dislocation. The thoracic cavity was opened, the descending aorta and inferior vena cava were severed, and EDTA buffer (130 mM NaCl, 5 mM KCl, 0.5 mM NaH_2_PO_4_, 10 mM HEPES, 10 mM glucose, 10 mM 2,3-butanedione monoxime, 5 mM EDTA, adjusted to pH 7.8) was injected into the right ventricle. The ascending aorta was clamped, and the heart was excised and transferred to a Petri dish containing EDTA buffer. A second injection was delivered into the left ventricle. The heart was then transferred to a perfusion dish containing perfusion buffer (same composition as EDTA buffer but with 1 mM MgCl_2_ in place of EDTA) and cleared of blood. The heart was placed on a 37°C heating pad and perfused with collagenase buffer (2.5 mg/mL collagenase type II, Worthington LS004177; 0.25 mg/mL protease from *Streptomyces griseus*, Sigma-Aldrich P5147, in perfusion buffer) at 1.2 mL/min for 20 minutes. Ventricles were dissected, transferred to a 60 mm Petri dish containing 5 mL collagenase buffer, and teased apart into approximately 1 mm × 1 mm pieces with forceps. Tissue was gently triturated 15 times with a transfer pipette and enzymatic digestion was terminated by addition of 10 mL stop solution (10% fetal bovine serum in perfusion buffer). The suspension was passed through a 100 μm cell strainer. Elapsed time from euthanasia to start of enzymatic digestion was under 5 minutes for every animal.

#### Fixation

CM-enriched suspensions were centrifuged at 20 × g for 5 minutes at room temperature, the supernatant was aspirated, and cells were fixed using the Evercode Cell Fixation kit (Parse Biosciences) according to the manufacturer’s instructions^38^. Fixed cells were resuspended in the supplied storage buffer, frozen at −80°C in a Mr. Frosty freezing container, and stored for less than 6 months prior to library preparation.

#### Single-cell library preparation and sequencing

Libraries were prepared using the Evercode Whole Transcriptome Mega kit (Parse Biosciences, chemistry v3) with samples distributed across one 96-well round-1 barcoding plate and six sublibraries. Reads were sequenced on an Illumina NovaSeq X Plus (paired-end, 64 bp + 58 bp) to a total depth of 20.9 billion reads.

#### Read processing and quality control

Reads were demultiplexed, aligned, and quantified using split-pipe (Parse Biosciences, v1.7.3) against a GRCm39 reference (Ensembl release 113)^39^. Cells were retained if they expressed at least 800 genes with upper bounds of 60,000 UMIs and 50% mitochondrial reads. Non-cardiomyocytes were identified by marker gene expression and removed, yielding the final CM populations used for atlas construction.

#### Animal details for jhu_p7_ctr and jhu_p14_ctr

Wild-type C57BL/6 mice were harvested at postnatal days 7 and 14 respectively. No viral injection or genetic modification was applied. The same Langendorff-free isolation and Parse SPLiT-seq library preparation protocol described above was followed.

#### Animal details for northwestern_ctr

Wild-type C57BL/6 mice were harvested at embryonic days 12.5, 14.5, and 16.5 and postnatal days 0 and 7. Animals received a subcutaneous injection of saline as a vehicle control. The same Langendorff-free isolation and Parse SPLiT-seq library preparation protocol described above was followed.

#### Animal details for Zureick_P14 external validation dataset

Homozygous B6J.129(B6N)-*Gt(ROSA)26Sor^tm1(CAG-cas9*,-EGFP)Fezh/^*J mice (Rosa26-LSL-Cas9-EGFP; The Jackson Laboratory, strain #026175) were harvested at postnatal day 14 (n=6; 3 male, 3 female) as part of a larger unpublished in vivo CRISPR screening experiment. Animals received a subcutaneous injection of AAV9 delivering Cre recombinase together with a sgRNA cassette at postnatal day 0. Only cardiomyocytes in which neither EGFP nor sgRNA transcripts were detected (untransduced cells, defined as zero eGFP counts and zero reads in the targeted CRISPR capture library) were retained for the analyses described here, yielding 98,163 cardiomyocytes across 6 biological replicates. These cells served as the external validation cohort for CMAC and were not used in any stage of atlas construction or model training

### Data preprocessing

#### Quality control

We sought to train on high quality cardiomyocytes, thus cell-level quality control was performed using metrics designed to enable standardized assessment across datasets generated using different experimental platforms and sequencing depths. For each cell, we quantified the proportion of reads mapping to the five most highly expressed genes (top5), mitochondrial genes (mito and mito_all), and ribosomal protein genes (ribo), together with library depth. To account for systematic differences in sequencing characteristics across studies and developmental stages, the proportion of reads assigned to the top five genes and sequencing depth were normalized within each dataset and developmental timepoint (top5_norm and depth_norm, respectively). Cardiomyocyte identity was independently assessed using SingleCellNet classification against the Tabula Muris reference^40,41^. Cells were retained for atlas construction when they satisfied the predefined quality-control criteria (top5_norm < 1.3 and depth_norm > −0.5) and were classified as cardiac muscle cells. This within-dataset, timepoint-normalized strategy was used to minimize the influence of technical differences across heterogeneous studies while avoiding the application of uniform absolute thresholds to datasets generated using distinct sequencing platforms and protocols.

#### Highly Variable Genes

Raw count matrices were harmonized to a common Ensembl gene universe using a master gene map. To identify genes exhibiting robust transcriptional variation across cardiomyocyte development while limiting the influence of differences in dataset size and developmental-stage representation, counts were normalized to 10,000 per cell and log1p-transformed. Gene expression was first averaged within each dataset at each developmental timepoint and subsequently averaged across datasets representing the same biological age, thereby assigning equal weight to each dataset within a developmental stage. Embryonic and postnatal timepoints were converted to a continuous biological-age scale, and genes were ranked according to the variance of their mean expression across developmental ages. Mitochondrial, ribosomal, and pseudogene features were excluded prior to variance ranking, and the top 10% of genes by developmental variance were retained. This procedure yielded 2,991 maturation-variable genes for atlas construction.

#### Dataset balancing

To prevent datasets or developmental stages with large numbers of cells from disproportionately influencing atlas construction and model training, cells were randomly downsampled to a maximum of 300 cells per dataset × biological-age combination. Groups containing fewer than 300 cells were retained in their entirety. This strategy preserved representation across datasets and developmental stages while limiting the contribution of highly sampled dataset–age combinations. Downsampling was performed without replacement using a fixed random seed to ensure reproducibility.

### Latent representation and atlas integration

#### scVI

A variational autoencoder implemented in scVI was trained on raw count data using a negative binomial likelihood with two hidden layers of 128 units and a 100-dimensional latent space.^16^ The dimensionality of the scVI latent representation was empirically optimized by evaluating latent spaces ranging from 30 to 150 dimensions for downstream biological-age prediction. Model performance was assessed using mean absolute error (MAE), Pearson and Spearman correlations between predicted and chronological age. A 100-dimensional latent space was selected because it achieved the lowest MAE (5.65 days) and highest Pearson correlation (r = 0.77), while maintaining a high Spearman correlation (ρ = 0.82). Increasing the latent dimensionality beyond 100 did not improve overall predictive performance, supporting 100 dimensions as an appropriate balance between representation capacity and generalization.Each dataset was assigned a unique batch label to account for platform and laboratory effects.

#### Integration metrics

Integration quality was evaluated using complementary metrics designed to assess both removal of dataset-specific structure and preservation of developmental organization within the scVI latent space. Dataset mixing was quantified using the dataset silhouette score (−0.076), cross-dataset nearest-neighbor fraction (55.4%), and integration Local Inverse Simpson’s Index (iLISI; median = 2.527). iLISI quantifies the effective diversity of dataset labels within local neighborhoods, with values above 1 indicating mixing of cells from multiple datasets. Preservation of developmental structure was assessed by the Spearman correlation between biological age and local neighborhood age (ρ = 0.893) and the median absolute age difference between neighboring cells (2.77 days). Together, these complementary metrics were used to evaluate whether integration reduced study-specific structure while retaining the developmental organization required for maturation modeling. UMAP visualization was generated from the scVI latent representation using 30 nearest neighbors.

### Developmental trajectory analysis

For each developmental timepoint, the mean scVI latent vector was calculated across all cells assigned to that stage. Cosine distance, which measures the angle between transcriptional program vectors independent of vector magnitude, was computed between consecutive timepoint centroids ordered chronologically from E7.75 to P28. Cosine distances were cumulatively summed along the ordered trajectory and normalized to a 0 to 1 scale, where 0 represents the earliest cardiac progenitor state, embryonic day 7.75 (e7.75) and 1.0 represents completed postnatal maturation, post-natal day 28(p28). Equal increments on this scale correspond to equal amounts of transcriptional change, providing a biologically calibrated description of developmental progression independent of chronological time.

### Chronological age prediction

Chronological age was predicted from the 100-dimensional scVI latent representation using XGBoost regression.^17^ Ages were modeled on the logarithmic scale and predictions were back-transformed by exponentiation to return age in days post-conception. Training observations were weighted inversely by dataset × developmental stage group size to reduce overrepresentation of densely sampled timepoints. Hyperparameters were selected via nested cross-validation using GroupKFold with 5 outer folds and 53 dataset × developmental stage groups, evaluating 48 parameter combinations per fold. Consensus parameters were identified from the most frequently selected values across all folds: learning_rate=0.05, n_estimators=600, max_depth=3, min_child_weight=1, subsample=0.8, colsample_bytree=0.8,reg_lambda=1.0, reg_alpha=0.0. Model performance was compared against a Ridge regression baseline and a PCA 100-dimensional + XGBoost approach using the same cross-validation design.

### Cross-dataset validation

Generalization was assessed using leave-one-dataset-out (LOGO) cross-validation across all 18 datasets. For each fold, the hyperparameter-selected XGBoost model was trained on 17 datasets with fixed consensus parameters and evaluated on the held-out dataset. Predictions were aggregated at the (dataset × developmental stage) level by computing mean predicted age per group, and performance was quantified using mean absolute error (MAE), Pearson correlation, and Spearman correlation between observed and predicted age across the 53 dataset × stage groups. Datasets covering only a single developmental timepoint contribute to MAE but not to correlation metrics, which require variance in the true age variable.

### PGC1α/β knockout validation

CMAC was applied to an independent PGC1α/β double-knockout cardiomyocyte dataset spanning postnatal days 7, 14, 21, and 28.^32^ Knockout cells were identified by RFP expression (RFP^+^) and wild-type controls by absence of RFP expression (RFP^−^). Both populations were projected into the CMAC latent space via scVI reference mapping using load_query_data with frozen encoder weights, and chronological age was predicted using the trained XGBoost model.^28^ Predicted age deficits were computed as the difference between median predicted age of RFP^+^ knockout and RFP^−^ control cells at each timepoint. As individual cell-level replication structure could not be confirmed for this dataset, statistical significance testing was not performed and effect sizes are reported. Deficits were interpreted relative to the CMAC median cross-validation error of 3.27 days, with deficits exceeding this threshold considered to reflect biological signal above model prediction uncertainty

## Data Availability

All public datasets used for atlas construction are available at accessions listed in Supplemental Table 1. Three in-house datasets (jhu_p7_ctr, jhu_p14_ctr, northwestern_ctr) and one external validation dataset (Zureick_P14) were generated in this study and have not been previously published; processed count matrices will be deposited to the Gene Expression Omnibus upon publication and are available from the corresponding author upon reasonable request in the interim. The trained CMAC model weights, including the scVI atlas model and XGBoost regressor, will be made publicly available upon publication. Analysis code will be deposited to GitHub upon publication. All data and code are available from the corresponding author upon reasonable request

## Code Availability

All analysis code used to construct the CMAC atlas, train the scVI and XGBoost models, perform cross-validation, and generate figures will be deposited to a public GitHub repository upon publication. The repository will include and a query projection pipeline enabling CMAC scoring of new cardiomyocyte datasets. Code is available from the corresponding author upon reasonable request prior to publication.

## Acknowledgement

This research was supported by the National Institutes of Health/National Heart, Lung, and Blood Institute (NIH/NHLBI) grants R01HL156947 and R01HL171205. We thank the members of the Kwon laboratory for helpful discussions.

## Author Contribution

K.Q. and S.M.: Conceptualization. K.Q., E.A., E.K., N.Z., E.C., and D.S.: Data Curation. K.Q.: Formal Analysis. N.Z. and K.Q.: Writing of original Draft. G.S.O., P.C., and C.K.: Supervision. C.K.: Funding Acquisition. C.K.: Review and Editing

## References

1. Li, B., Liu, X., Murphy, S., Tampakakis, E., and Kwon, C. (2026). Decoding Cardiac Development and Maturation at Single-Cell and Spatial Transcriptomic Resolution. Circulation research 139. 10.1161/CIRCRESAHA.125.327473.

2. Murphy, S.A., Chen, E.Z., Tung, L., Boheler, K.R., and Kwon, C. (2021). Maturing heart muscle cells: Mechanisms and transcriptomic insights. Semin Cell Dev Biol 119, 49–60. 10.1016/j.semcdb.2021.04.019.

3. Hershberger, R.E., Morales, A., and Siegfried, J.D. (2010). Clinical and genetic issues in dilated cardiomyopathy: a review for genetics professionals. Genet Med 12, 655–667. 10.1097/GIM.0b013e3181f2481f.

4. Seidman, J.G., and Seidman, C. (2001). The Genetic Basis for Cardiomyopathy: from Mutation Identification to Mechanistic Paradigms. Cell 104, 557–567. 10.1016/S0092-8674(01)00242-2.

5. Lundy, S.D., Zhu, W.-Z., Regnier, M., and Laflamme, M.A. (2013). Structural and functional maturation of cardiomyocytes derived from human pluripotent stem cells. Stem Cells Dev 22, 1991–2002. 10.1089/scd.2012.0490.

6. Mummery, C.L., Zhang, J., Ng, E.S., Elliott, D.A., Elefanty, A.G., and Kamp, T.J. (2012). Differentiation of human embryonic stem cells and induced pluripotent stem cells to cardiomyocytes: a methods overview. Circ Res 111, 344–358. 10.1161/CIRCRESAHA.110.227512.

7. Guo, Y., and Pu, W.T. (2020). Cardiomyocyte Maturation. Circulation Research 126, 1086–1106. 10.1161/CIRCRESAHA.119.315862.

8. Quansah, K., Murphy, S., Kwon, E., Anderson, E., Tichnell, C., Murray, B., Gaine, S., Gasperetti, A., James, C., Calkins, H., et al. (2026). Machine Learning Models Enhance Detection of Arrhythmogenic Right Ventricular Cardiomyopathy. Mach. Learn.: Health. 10.1088/3049-477X/ae451a.

9. Quansah, K., Anderson, E., Kwon, E., and Kwon, C. (2026). Machine Learning in Nonischemic Cardiomyopathy: Phenotyping, Mechanism Discovery, and Clinical Applications. Cardiol Rev. 10.1097/CRD.0000000000001281.

10. Chanthra, N., Abe, T., Miyamoto, M., Sekiguchi, K., Kwon, C., Hanazono, Y., and Uosaki, H. (2020). A Novel Fluorescent Reporter System Identifies Laminin-511/521 as Potent Regulators of Cardiomyocyte Maturation. Sci Rep 10, 4249. 10.1038/s41598-020-61163-3.

11. DeLaughter, D.M., Bick, A.G., Wakimoto, H., McKean, D., Gorham, J.M., Kathiriya, I.S., Hinson, J.T., Homsy, J., Gray, J., Pu, W., et al. (2016). Single-Cell Resolution of Temporal Gene Expression during Heart Development. Dev Cell 39, 480–490. 10.1016/j.devcel.2016.10.001.

12. Trapnell, C., Cacchiarelli, D., Grimsby, J., Pokharel, P., Li, S., Morse, M., Lennon, N.J., Livak, K.J., Mikkelsen, T.S., and Rinn, J.L. (2014). The dynamics and regulators of cell fate decisions are revealed by pseudotemporal ordering of single cells. Nat Biotechnol 32, 381–386. 10.1038/nbt.2859.

13. Chen, E.Z., Kannan, S., Murphy, S., Farid, M., and Kwon, C. (2024). Protocol for quantifying stem-cell-derived cardiomyocyte maturity using transcriptomic entropy score. STAR Protoc 5, 103083. 10.1016/j.xpro.2024.103083.

14. Tung, P.-Y., Blischak, J.D., Hsiao, C.J., Knowles, D.A., Burnett, J.E., Pritchard, J.K., and Gilad, Y. (2017). Batch effects and the effective design of single-cell gene expression studies. Sci Rep 7, 39921. 10.1038/srep39921.

15. Luecken, M.D., and Theis, F.J. (2019). Current best practices in single-cell RNA-seq analysis: a tutorial. Mol Syst Biol 15, e8746. 10.15252/msb.20188746.

16. Lopez, R., Regier, J., Cole, M.B., Jordan, M.I., and Yosef, N. (2018). Deep generative modeling for single-cell transcriptomics. Nat Methods 15, 1053–1058. 10.1038/s41592-018-0229-2.

17. Chen, T., and Guestrin, C. (2016). XGBoost: A Scalable Tree Boosting System. In Proceedings of the 22nd ACM SIGKDD International Conference on Knowledge Discovery and Data Mining, pp. 785–794. 10.1145/2939672.2939785.

18. De Soysa, T.Y., Ranade, S.S., Okawa, S., Ravichandran, S., Huang, Y., Salunga, H.T., Schricker, A., Del Sol, A., Gifford, C.A., and Srivastava, D. (2019). Single-cell analysis of cardiogenesis reveals basis for organ-level developmental defects. Nature 572, 120–124. 10.1038/s41586-019-1414-x.

19. Kannan, S., Farid, M., Lin, B.L., Miyamoto, M., and Kwon, C. (2021). Transcriptomic entropy benchmarks stem cell-derived cardiomyocyte maturation against endogenous tissue at single cell level. PLOS Computational Biology 17, e1009305. 10.1371/journal.pcbi.1009305.

20. Goodyer, W.R., Beyersdorf, B.M., Paik, D.T., Tian, L., Li, G., Buikema, J.W., Chirikian, O., Choi, S., Venkatraman, S., Adams, E.L., et al. (2019). Transcriptomic Profiling of the Developing Cardiac Conduction System at Single-Cell Resolution. Circulation Research 125, 379–397. 10.1161/CIRCRESAHA.118.314578.

21. Hill, M.C., Kadow, Z.A., Li, L., Tran, T.T., Wythe, J.D., and Martin, J.F. (2019). A cellular atlas of Pitx2-dependent cardiac development. Development 146, dev180398. 10.1242/dev.180398.

22. Pijuan-Sala, B., Griffiths, J.A., Guibentif, C., Hiscock, T.W., Jawaid, W., Calero-Nieto, F.J., Mulas, C., Ibarra-Soria, X., Tyser, R.C.V., Ho, D.L.L., et al. (2019). A single-cell molecular map of mouse gastrulation and early organogenesis. Nature 566, 490–495. 10.1038/s41586-019-0933-9.

23. Xiong, H., Luo, Y., Yue, Y., Zhang, J., Ai, S., Li, X., Wang, X., Zhang, Y.-L., Wei, Y., Li, H.-H., et al. (2019). Single-Cell Transcriptomics Reveals Chemotaxis-Mediated Intraorgan Crosstalk During Cardiogenesis. Circulation Research 125, 398–410. 10.1161/CIRCRESAHA.119.315243.

24. Xiao, Y., Hill, M.C., Zhang, M., Martin, T.J., Morikawa, Y., Wang, S., Moise, A.R., Wythe, J.D., and Martin, J.F. (2018). Hippo Signaling Plays an Essential Role in Cell State Transitions during Cardiac Fibroblast Development. Dev Cell 45, 153–169.e6. 10.1016/j.devcel.2018.03.019.

25. Feulner, L., Wünnemann, F., Liang, J., Hofmann, P., Hitz, M.-P., Schapiro, D., Leclerc, S., Vliet, P.P. van, and Andelfinger, G. (2024). Transcriptional dynamics of the murine heart during perinatal development at single-cell resolution. Preprint at bioRxiv, 10.1101/2024.03.05.583423 https://doi.org/10.1101/2024.03.05.583423.

26. Wang, Y., Yao, F., Wang, L., Li, Z., Ren, Z., Li, D., Zhang, M., Han, L., Wang, S., Zhou, B., et al. (2020). Single-cell analysis of murine fibroblasts identifies neonatal to adult switching that regulates cardiomyocyte maturation. Nat Commun 11, 2585. 10.1038/s41467-020-16204-w.

27. Dong, J., Hu, Y., Fan, X., Wu, X., Mao, Y., Hu, B., Guo, H., Wen, L., and Tang, F. (2018). Single-cell RNA-seq analysis unveils a prevalent epithelial/mesenchymal hybrid state during mouse organogenesis. Genome Biol 19, 31. 10.1186/s13059-018-1416-2.

28. Lotfollahi, M., Naghipourfar, M., Luecken, M.D., Khajavi, M., Büttner, M., Wagenstetter, M., Avsec, Ž., Gayoso, A., Yosef, N., Interlandi, M., et al. (2022). Mapping single-cell data to reference atlases by transfer learning. Nat Biotechnol 40, 121–130. 10.1038/s41587-021-01001-7.

29. Beisaw, A., and Wu, C.-C. (2024). Cardiomyocyte maturation and its reversal during cardiac regeneration. Developmental Dynamics 253, 8–27. 10.1002/dvdy.557.

30. Li, Z., Yao, F., Yu, P., Li, D., Zhang, M., Mao, L., Shen, X., Ren, Z., Wang, L., and Zhou, B. (2022). Postnatal state transition of cardiomyocyte as a primary step in heart maturation. protein. cell. 13, 842–862. 10.1007/s13238-022-00908-4.

31. Lopaschuk, G.D., Ussher, J.R., Folmes, C.D.L., Jaswal, J.S., and Stanley, W.C. (2010). Myocardial fatty acid metabolism in health and disease. Physiol Rev 90, 207–258. 10.1152/physrev.00015.2009.

32. Murphy, S.A., Miyamoto, M., Kervadec, A., Kannan, S., Tampakakis, E., Kambhampati, S., Lin, B.L., Paek, S., Andersen, P., Lee, D.-I., et al. (2021). PGC1/PPAR drive cardiomyocyte maturation at single cell level via YAP1 and SF3B2. Nat Commun 12, 1648. 10.1038/s41467-021-21957-z.

33. Garbern, J.C., and Lee, R.T. (2021). Mitochondria and metabolic transitions in cardiomyocytes: lessons from development for stem cell-derived cardiomyocytes. Stem Cell Res Ther 12, 177. 10.1186/s13287-021-02252-6.

34. Tan, C.M.J., and Lewandowski, A.J. (2019). The Transitional Heart: From Early Embryonic and Fetal Development to Neonatal Life. Fetal Diagn Ther 47, 373–386. 10.1159/000501906.

35. Lopaschuk, G.D., and Jaswal, J.S. (2010). Energy metabolic phenotype of the cardiomyocyte during development, differentiation, and postnatal maturation. J Cardiovasc Pharmacol 56, 130–140. 10.1097/FJC.0b013e3181e74a14.

36. Friedman, C.E., Nguyen, Q., Lukowski, S.W., Helfer, A., Chiu, H.S., Miklas, J., Levy, S., Suo, S., Han, J.-D.J., Osteil, P., et al. (2018). Single-Cell Transcriptomic Analysis of Cardiac Differentiation from Human PSCs Reveals HOPX-Dependent Cardiomyocyte Maturation. Cell Stem Cell 23, 586–598.e8. 10.1016/j.stem.2018.09.009.

37. Ackers-Johnson, M., Li, P.Y., Holmes, A.P., O’Brien, S.-M., Pavlovic, D., and Foo, R.S. (2016). A Simplified, Langendorff-Free Method for Concomitant Isolation of Viable Cardiac Myocytes and Nonmyocytes From the Adult Mouse Heart. Circ Res 119, 909–920. 10.1161/CIRCRESAHA.116.309202.

38. Parse Biosciences (2024). Split-pipe Pipeline. Version 1.7.3 (Parse Biosciences).

39. Ensembl (2024). Mouse Genome Assembly GRCm39, Ensembl Release 113. https://ensembl.org https://ensembl.org.

40. Schaum, N., Karkanias, J., Neff, N.F., May, A.P., Quake, S.R., Wyss-Coray, T., Darmanis, S., Batson, J., Botvinnik, O., Chen, M.B., et al. (2018). Single-cell transcriptomics of 20 mouse organs creates a Tabula Muris. Nature 562, 367–372. 10.1038/s41586-018-0590-4.

41. Tan, Y., and Cahan, P. (2019). SingleCellNet: a computational tool to classify single cell RNA-Seq data across platforms and across species. Cell Syst 9, 207–213.e2. 10.1016/j.cels.2019.06.004.

