## Supplemental Table 1 for "CMAC: A deep learning framework for absolute cardiomyocyte transcriptional age estimation"

| Dataset | Author | Isolation | Sequencing | Mapping | Datatype | TImepoints | Accession_number | Species | sample_type |
|---|---|---|---|---|---|---|---|---|---|
| chromium_data | 10x Chromium Demo | 10x Chromium V3 | 10x Chromium | CellRanger | UMIs | e18 | 10x website | mouse | in vivo |
| desoysa_data | De Soysa | 10x Chromium V2 | 10x Chromium | CellRanger | UMIs | e7.75, e8.25, e9.25 | GSE126128 | mouse | in vivo |
| dong_data | Dong | Picked | 8bp-STRT-seq | Kallisto/Bustools | UMIs | e9.5, e10.5, e11.5 | GSE87038 | mouse | in vivo |
| goodyer_avn_data | Goodyer AVN | 10x Chromium V2 | 10x Chromium | CellRanger | UMIs | e16.5 | GSE132658 | mouse | in vivo |
| goodyer_pf_data | Goodyer PF | 10x Chromium V2 | 10x Chromium | CellRanger | UMIs | e16.5 | GSE132658 | mouse | in vivo |
| hill10_data | Hill e10.5 | 10x Chromium V2 | 10x Chromium | CellRanger | UMIs | e10.5 | GSE131181 | mouse | in vivo |
| hill13_data | Hill e13.5 | 10x Chromium V2 | 10x Chromium | CellRanger | UMIs | e13.5 | GSE131181 | mouse | in vivo |
| jhu_p7_ctr | Suh, Chen | Langendorf-free, Ackers-Johnson | Parse split-seq | Trailmaker/split-pipe | UMIs | p7 | unpublished | mouse | in vivo |
| jhu_p14_ctr | Suh, Chen | Langendorf-free, Ackers-Johnson | Parse split-seq | Trailmaker/split-pipe | UMIs | p14 | unpublished | mouse | in vivo |
| kannan_ref_data | Kannan Reference | LP-FACS | mcSCRB-seq | zUMIs | UMIs | e18, e14, p0, p4, p14, p22, p28, p35, p56, p18, p11, p8 | GSE147807 | mouse | in vivo |
| li10_data | Li 10x | 10x Chromium V2 | 10x Chromium | CellRanger | UMIs | e10.5 | GSE76118 | mouse | in vivo |
| murphy_data | Murphy | LP-FACS | SCRB-seq | zUMIs | UMIs | p0, p7, p28, p21, p14 | GSE165917 | mouse | in vivo |
| northwestern_ctr | Suh | | Parse split-seq | Trailmaker/split-pipe | UMIs | e12.5, e14.5, e16.5, P0, P7 | unpublished | mouse | in vivo |
| pijuansala_data | Pijuan-Sala, Griffiths, and Guibentif | 10x Chromium V1 | 10x Chromium | CellRanger | UMIs | e7.75, e8.0, e8.5, e8.25 | doi:10.1038/s41586-019-0933-9 | mouse | in vivo |
| wang_dev_data | Wang and Yao | iCell8 | Unknown | STAR/FeatureCounts | UMIs | p1, p4, p7, p14 | GSE122706 | mouse | in vivo |
| wunnemann_data | Wunnemann | Drop-seq | Drop-seq | Drop-seq Tools | UMIs | e14.5, e16.5, e18.5, p1, p4, p7 | GSE150817, GSE109247 | mouse | in vivo |
| xiao_data | Xiao | Drop-seq | Drop-seq | Drop-seq Tools | UMIs | e13.5, e14.5 | GSE100861 | mouse | in vivo |
| xiong_data | Xiong, Luo, Yue, and Zhang | Picked | 8bp-STRT-seq | Kallisto/Bustools | UMIs | e9.25, e8.25, e8.75 | GSE108963 | mouse | in vivo |
| zureick_P14 | Zureick, Chen | Langendorf-free, Ackers-Johnson | Parse split-seq | Trailmaker/split-pipe | UMIs | p14 | unpublished | mouse | in vivo |
